# SR-B1 uptake of HDL promotes prostate cancer proliferation and tumor progression

**DOI:** 10.1101/2020.02.24.963454

**Authors:** C. Alicia Traughber, Emmanuel Opoku, Gregory Brubaker, Jennifer Major, Hanxu Lu, Shuhui Wang Lorkowski, Chase Neumann, Aimalie Hardaway, Yoon-Mi Chung, Kailash Gulshan, Nima Sharifi, J. Mark Brown, Jonathan D. Smith

## Abstract

High density lipoprotein (HDL) metabolism, in part, is facilitated by scavenger receptor class B, type 1 (SR-B1) that mediates its uptake into cells. SR-B1 is upregulated in prostate cancer tissue. Here, we report that knockout (KO) of SR-B1 via CRISPR/Cas9 editing led to reduced HDL uptake into prostate cancer cells, and reduced their proliferation in response to HDL. *In vivo* studies using syngeneic SR-B1 wildtype (SR-B1^+/+^) and SR-B1 KO (SR-B1^−/−^) prostate cancer cells in WT and apolipoprotein-AI KO (apoA1-KO) C57BL/6J mice showed that WT hosts, containing higher levels of total and HDL-cholesterol, grew larger tumors than apoA1-KO hosts with lower levels of total and HDL-cholesterol. Furthermore, SR-B1^−/−^ prostate cancer cells formed smaller tumors in WT hosts, than SR-B1^+/+^ cells in same host model. Tumor volume data was overall similar to survival data. We conclude that tumoral SR-B1 KO reduced HDL-mediated increases in prostate cancer cell proliferation and disease progression.

---

Prostate cancer is the most common malignancy and second leading cause of cancer-related deaths among men in the United States (1). The association of HDL levels with prostate cancer risk has been inconsistent, with some studies showing a positive association (2,3) some showing an inverse association (4,5), and others showing no association (6,7). HDL biogenesis is mediated by ATP-binding cassette transporter A1 (ABCA1), which assembles cellular lipids with exogenous lipid-poor apolipoprotein A1 (apoA1) to generate nascent HDL (8). HDL uptake in tissues is facilitated by SR-B1 (9), which can also mediate bidirectional cholesterol transport between cells and HDL (10–13). SR-B1, encoded by the *SCARB1* gene, is highly expressed in the liver, and even more so in steroidogenic tissues, such as the adrenals, testes, and ovaries where it mediates cholesterol uptake, to promote cholesterol ester (CE) storage used for steroid hormone synthesis (14,15).

It has been reported that prostate cancer accumulates CE in lipid droplets that correlates with prostate cancer aggressiveness (16). This phenotype was attributed to PTEN deletion that ultimately resulted in upregulation of the LDL receptor (LDLR) and subsequent uptake of LDL cholesterol (16). SR-B1 is inducible by androgens in human hepatoma cells and primary monocyte macrophage (17). Moreover, reports suggest that androgens during puberty are responsible for lower HDL levels in men vs. women, most likely due to higher hepatic SR-B1 levels (18–21). A prior study found that SR-B1 is upregulated in high grade vs. low grade prostate cancer, and in metastatic vs. primary prostate cancer, while the LDLR was not altered in high grade or metastatic prostate cancer (22). Furthermore it was shown that that high vs. low SR-B1 expression in prostatectomy specimens was associated with decreased progression-free survival (22). In a small study, SR-B1 mRNA levels were significantly higher in prostate cancer tissue vs. matched normal prostate tissue (23). In our study, we examined the effect of HDL on prostate cancer cell growth, proliferation, and tumor progression. We found that HDL, in an SR-B1-dependent manner, promoted increased prostate cancer cell growth *in vitro*. In a syngeneic mouse model, a high HDL environment promoted tumor progression in an SR-B1-dependent manner. These results suggest that SR-B1 and HDL uptake promote prostate cancer progression and that inhibiting HDL uptake may be a viable target for decreasing disease burden.

## Results

### SCARB1 is upregulated in human prostate cancer

Previous studies have focused on lipoprotein receptors in prostate cancer, and how their changes may influence cholesterol transport in prostate cancer (16,22–24). Therefore, we evaluated expression of SR-B1, LDLR, and ABCA1 in RNA-seq data from a total of 52 paired prostate cancer and normal adjacent tissue from the TCGA PRAD dataset (25). Since the LDLR expression levels in normal prostate were not normally distributed, non-parametric statistics were used for all gene expression data. The median log_2_ expression levels of *SCARB1* mRNA in normal prostate and prostate cancer were 9.82 and 10.57, respectively, representing a 68% increase in SR-B1 mRNA in prostate cancer (p<0.0001, Fig. 1A). However, *LDLR* (Fig. 1B) in prostate cancer tumor tissue remained unchanged (p=0.28), whereas *ABCA1* expression (Fig. 1C) was significantly decreased by 23% in the tumor tissue, (p=0.014), in agreement with a previous report (24).

**Figure 1.**
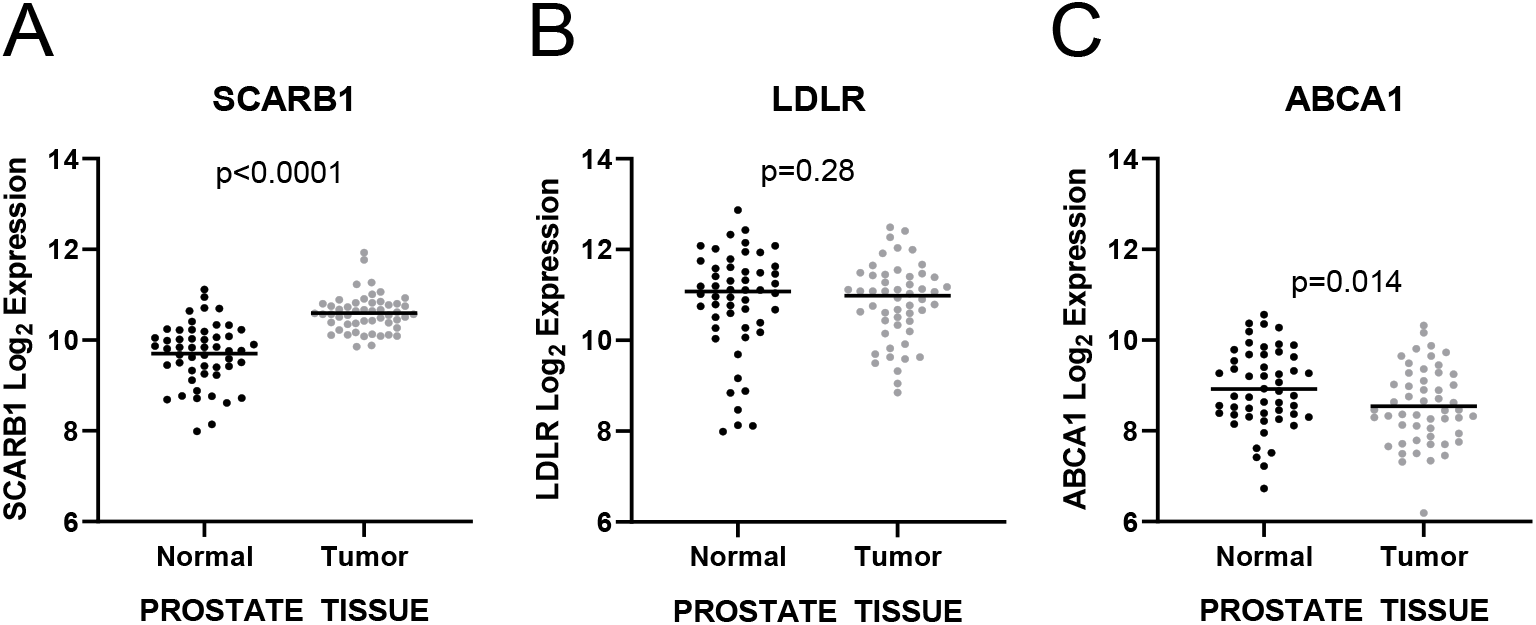
Expression of lipoprotein metabolism genes in normal adjacent and prostate tumor tissue. mRNA expression of A) *SCARB1*, B) *LDLR*, and C) *ABCA1* analyzed from TCGA PRAD RNA sequencing data (N=52 paired samples; median values shown; Wilcoxon paired non-parametric signed rank test with p-values displayed).

### HDL increased cell proliferation and cholesterol levels in prostate cancer cells

Since other studies (22,23) and our analysis of the TCGA data set showed upregulation of SR-B1 in prostate cancer (Fig. 1), we hypothesized that HDL may drive an increase of total cholesterol levels, similarly to what was previously illustrated for LDL (16). WT Human DU145 and mouse TRAMP-C2, expressing SR-B1 (SR-B1^+/+^), prostate cancer cells were incubated with 200 μg/ml HDL for 2 days leading to 48% (p<0.01) and 15% (p<0.05) increases in total cellular cholesterol levels (Fig 2A, B). We next determined if HDL could promote proliferation of prostate cancer cells by evaluating the impact on cell number. DU145 and TRAMP-C2 cells were treated with or without 300 μg/ml HDL in LPDS for 4 days. HDL induced 29% (p=0.0061) and 68% (p<0.0001) increases in cell number in DU145 and TRAMP-C2 cells, respectively (Fig 3A, B). To determine if this increase in cell number was due to increased cell proliferation, we performed a PKH26 dye-dilution assay in absence or presence of 300 μg/ml HDL. Flow cytometry analysis demonstrated that median fluorescence intensity (MFI) were significantly lower in HDL treated vs. untreated cells indicating more rounds of proliferation in both DU145 (26% decrease, p<0.001) and TRAMP-C2 (17% decrease, p<0.001) prostate cancer cells (Fig 3C, D).

**Figure 2.**
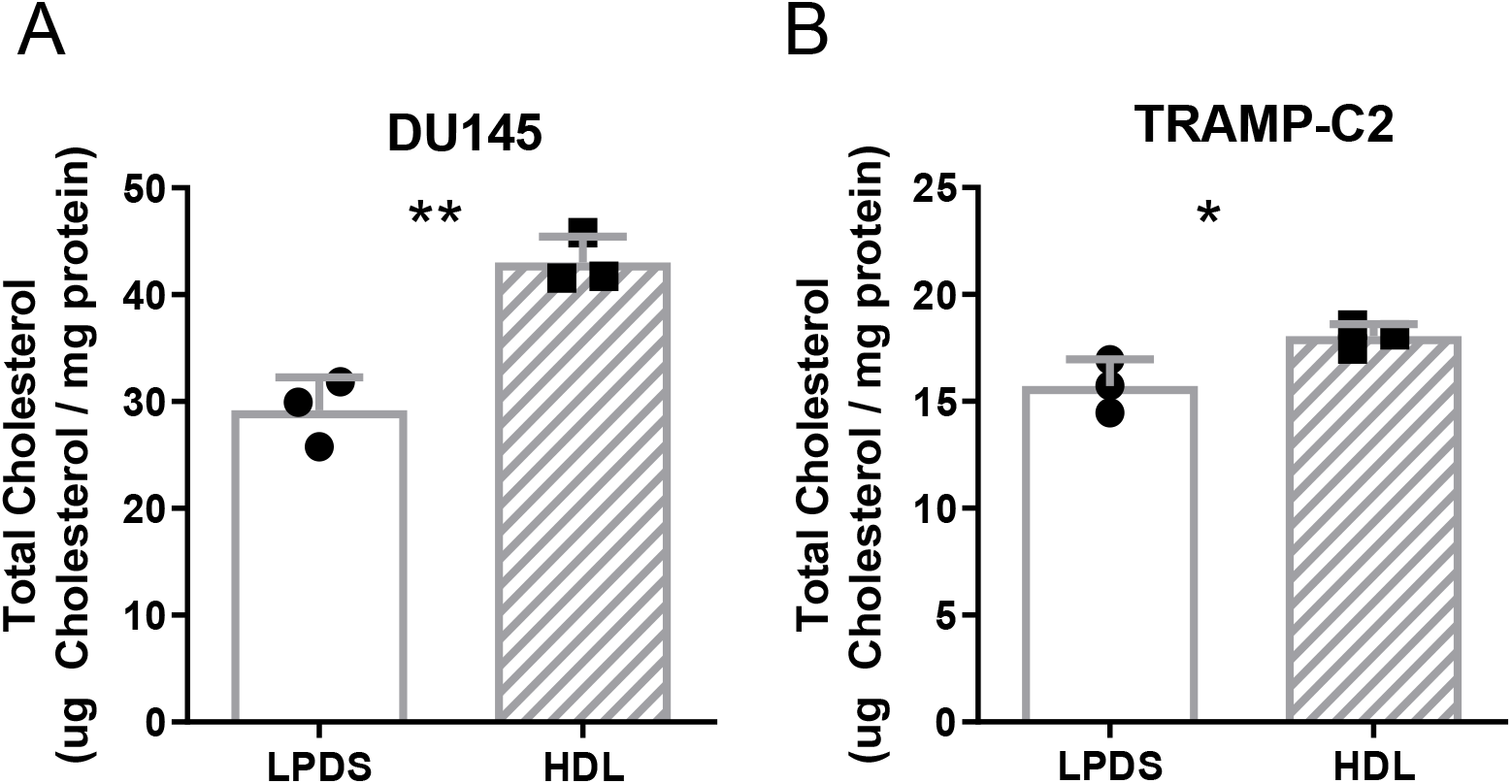
HDL effects on total cholesterol levels in prostate cancer cells. A) human DU145 and B) mouse TRAMP-C2 cells were incubated with ± 200 μg/ml HDL for 2 days in LPDS and total cholesterol levels normalized to cell protein determined (N=3; mean ± SD; *, p<0.05; **, p<0.01, by t-test).

**Figure 3.**
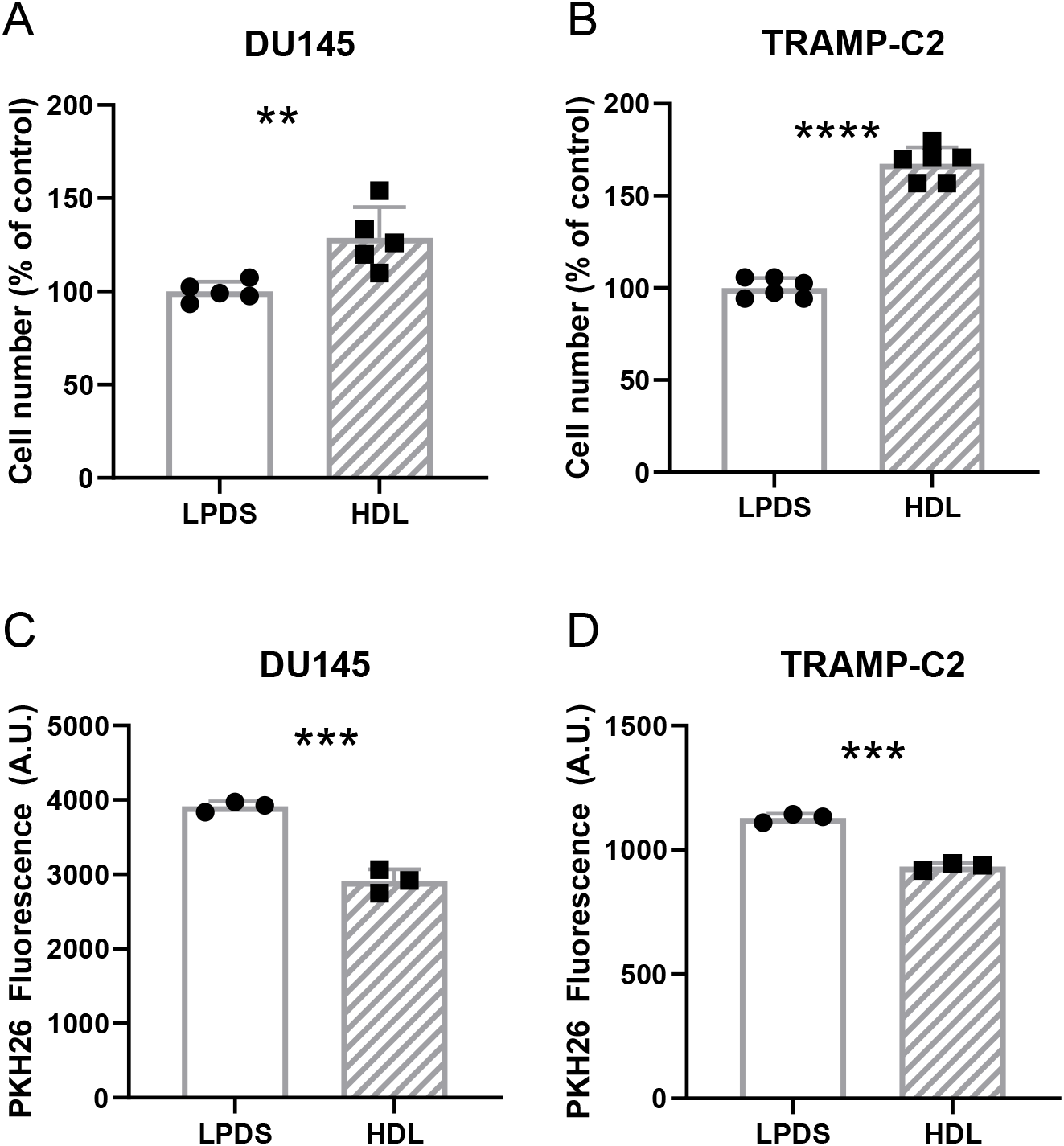
HDL effects on cell accumulation and proliferation of prostate cancer cells. A) human DU145 (N= 5 over 2 independent experiments) and B) mouse TRAMP-C2 (N=6 over 3 independent experiments) cells were treated ± 300 μg/ml HDL in LPDS media for 4 days and the final cell counts were normalized to the LPDS control. C, D) human DU145 and mouse TRAMP-C2 cells were labeled with the dye PKH26 and treated with ± 300 μg/ml HDL in LPDS media for 5 days and final dye dilution was analyzed by flow cytometry. Values are expressed as the mean ± SD; **p<0.01; ***, p<0.001. ****p<0.0001 by t-test.

### SR-B1 is required for HDL-mediated prostate cancer growth in vitro

Due to the response of prostate cancer cells to HDL, we next determined if these HDL effects were mediated by SR-B1. Therefore, we knocked out SR-B1 in both the human and mouse prostate cancer cell lines using CRISPR/Cas9 targeting exon 4, an early coding exon of the *SCARB1* gene, to generate cell lines with complete knockout of SR-B1 expression. Western blot demonstrated successful SR-B1 KO clones (SR-B1^−/−^) for both DU145 and TRAMP-C2 (Fig. 4A). Although HDL increased the relative cell number for both DU145 SR-B1^+/+^ (59.3% increase, p<0.0003) and SR-B1^−/−^ cells (23.4% increase, p=0.011), the increase in cell accumulation in response to a 4-day 200 μg/ml HDL treatment was significantly attenuated upon knockout of SR-B1 (Fig. 4B, p<0.0003, % control cell number in HDL-treated SR-B1^+/+^ vs. SR-B1^−/−^). We isolated three independent SR-B1^−/−^ clonally-derived cell lines from edited TRAMP-C2 cells, and their cell accumulation in LPDS was evaluated, showing that the SR-B1^−/−^ #5 line accumulated the least cells (p=0.02 vs. TRAMP-C2 SR-B1^+/+^), the SR-B1- #10 line accumulated the most cells (NS vs. TRAMP-C2 SR-B1^+/+^), and the SR-B1^−/−^ #17 line was most similar in cell number to SR-B1^+/+^ TRAMP-C2 cells (NS, Fig. 4C). The response of these cell lines to HDL was evaluated, normalized to the LPDS control for each of these lines. HDL significantly increased cell accumulation in WT cells (98% increase p=0.011), but not in any of the three SR-B1^−/−^ lines (Fig. 4D). Thus, we chose to further utilize and characterize HDL treatment on cellular processes in the SR-B1^−/−^ #17 cell line, as its basal growth levels in LPDS were the most similar to that of the TRAMP-C2 SR-B1^+/+^ cells (Fig. 4C).

**Figure 4.**
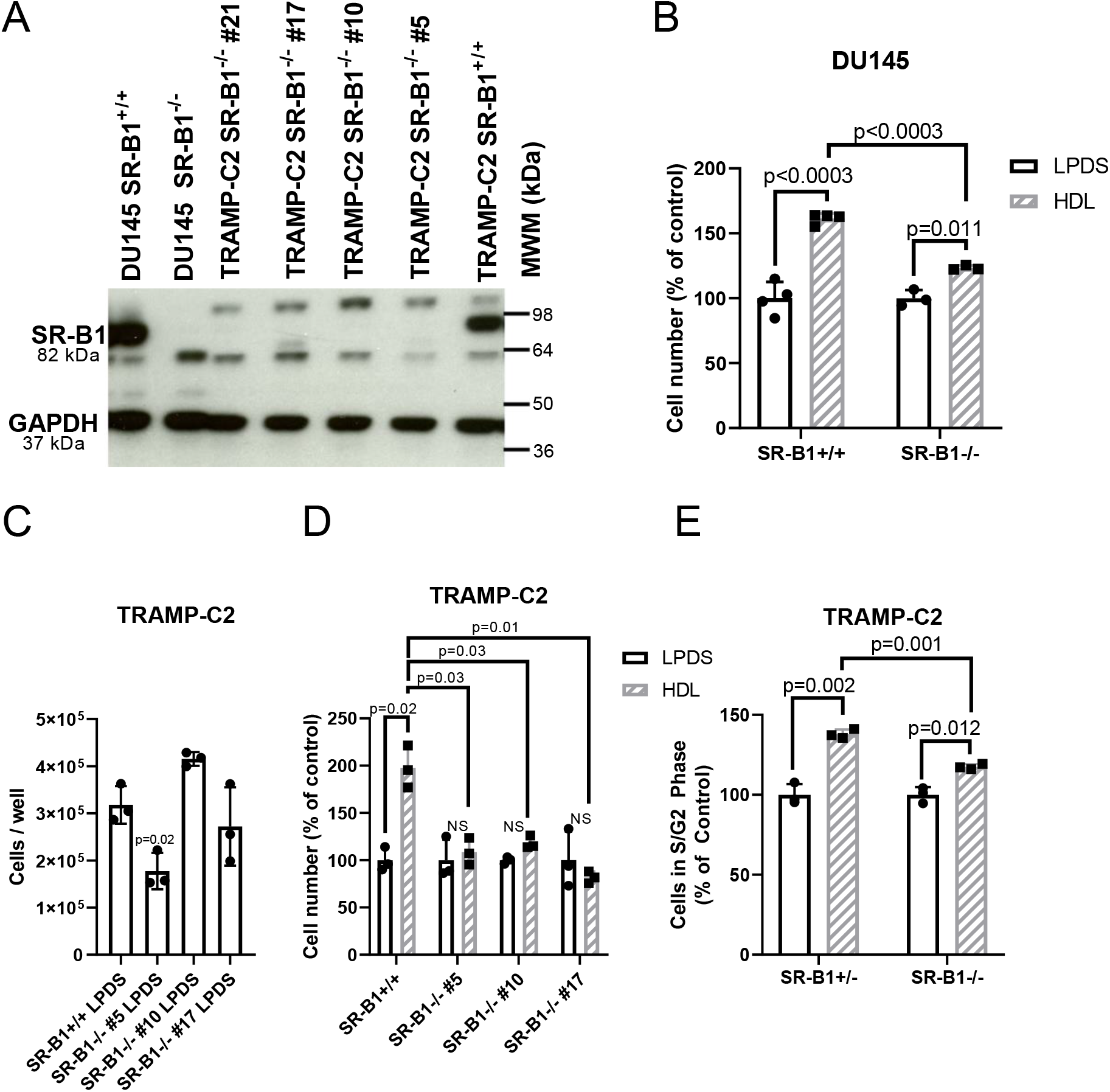
HDL effects in WT and SR-B1 KO cells. A) Western blot for SR-B1 in WT and SR-B1 KO DU145 and TRAMP-C2 cells. B) Cell accumulation assay for DU145 WT and SR-B1 KO cells incubated ± 300 μg/ml HDL in LPDS for 3 days (N=3-4; mean ± SD; t-test with Bonferroni correction for 3 tests; p-values displayed). C) Cell number of SR-B1^+/+^ TRAMP-C2 cells and three independent SR-B1^−/−^ clonally derived cell lines after incubation in 10% LPDS for 3 days (N=3; mean ± SD; ANOVA with Dunnett’s post-test comparing to SR-B1^+/+^ cells; p-value displayed). D) Cell accumulation in TRAMP-C2 SR-B1^+/+^ and three SR-B1^−/−^ clones incubated ± 200 μg/ml HDL for 3 days normalized to each cell lines LPDS control (N=3; mean ± SD; t-test with Bonferroni correction for 7 tests (4 tests ± HDL for each line and 3 tests of HDL treated SR-B1^+/+^ vs. SR-B1^−/−^ clones), significant p-values displayed). E) Cell cycle analysis in TRAMP-C2 SR-B1^+/+^ and SR-B1^−/−^ cells treated ± 300 μg/ml HDL in LPDS media for 1 day (% of cells in S+G2 phases; N=3; mean ± SD; t-test with Bonferroni correction for 3 tests, p-values displayed).

We next investigated if absence of SR-B1 impacts cell cycling upon HDL treatment by treating TRAMP-C2 SR-B1^+/+^ and SR-B1^−/−^ cells with or without 300 μg/ml HDL in LPDS for 1 day, then assessing the fraction of cycling cells in G_2_+S phase via propidium iodide content. We demonstrated that HDL increased the proportion of SR-B1^+/+^ cells cycling by 38% vs. the LPDS control (p=0.002), whereas in SR-B1^−/−^ cells there was only a 17.5% increase cycling cells in response to HDL (p=0.012, Fig. 4E). The HDL effect on cell cycling was significantly greater in SR-B1^+/+^ vs. SR-B1^−/−^ cells (p=0.001, Fig. 4E).

To confirm that HDL-uptake was impacted upon KO of SR-B1, we incubated TRAMP-C2 SR-B1^+/+^ and SR-B1^−/−^ cells with Alexa568-HDL. Fluorescent microscopy showed that SR-B1^+/+^ cells took up more Alexa568-HDL as compared to SR-B1^−/−^ cells (Fig. 5A). This was further confirmed by flow cytometry, where the cells were treated with 20 μg/ml Alexa568-HDL with or without 2 mg/ml unlabeled-HDL competitor. The cellular uptake of Alexa568-HDL was determined by median fluorescence intensity (MFI) showing that the SR-B1^−/−^ cells had reduced total and specific Alexa568-HDL uptake compared to the SR-B1^+/+^ cells by 44.1% (p=0.002) and 59.6%(p=0.007), respectively (Fig. 5B). To investigate if cholesterol is involved in the HDL effect on cell accumulation, we treated SR-B1^+/+^ or SR-B1^−/−^ cells with or without 1 μM lovastatin to reduce endogenous cholesterol biosynthesis, which significantly decreased cell accumulation in both cell lines (p<0.05, Fig. 5C). HDL treatment for 4 days added to the lovastatin significantly rescued the cell accumulation only in the SR-B1^+/+^ cells (p<0.05 vs. lovastatin alone, Fig. 5C). This suggests that HDL provides cholesterol, in an SR-B1-dependent manner, to help cells grow when *de novo* cholesterol was reduced by statin treatment (Fig 5C).

**Figure 5.**
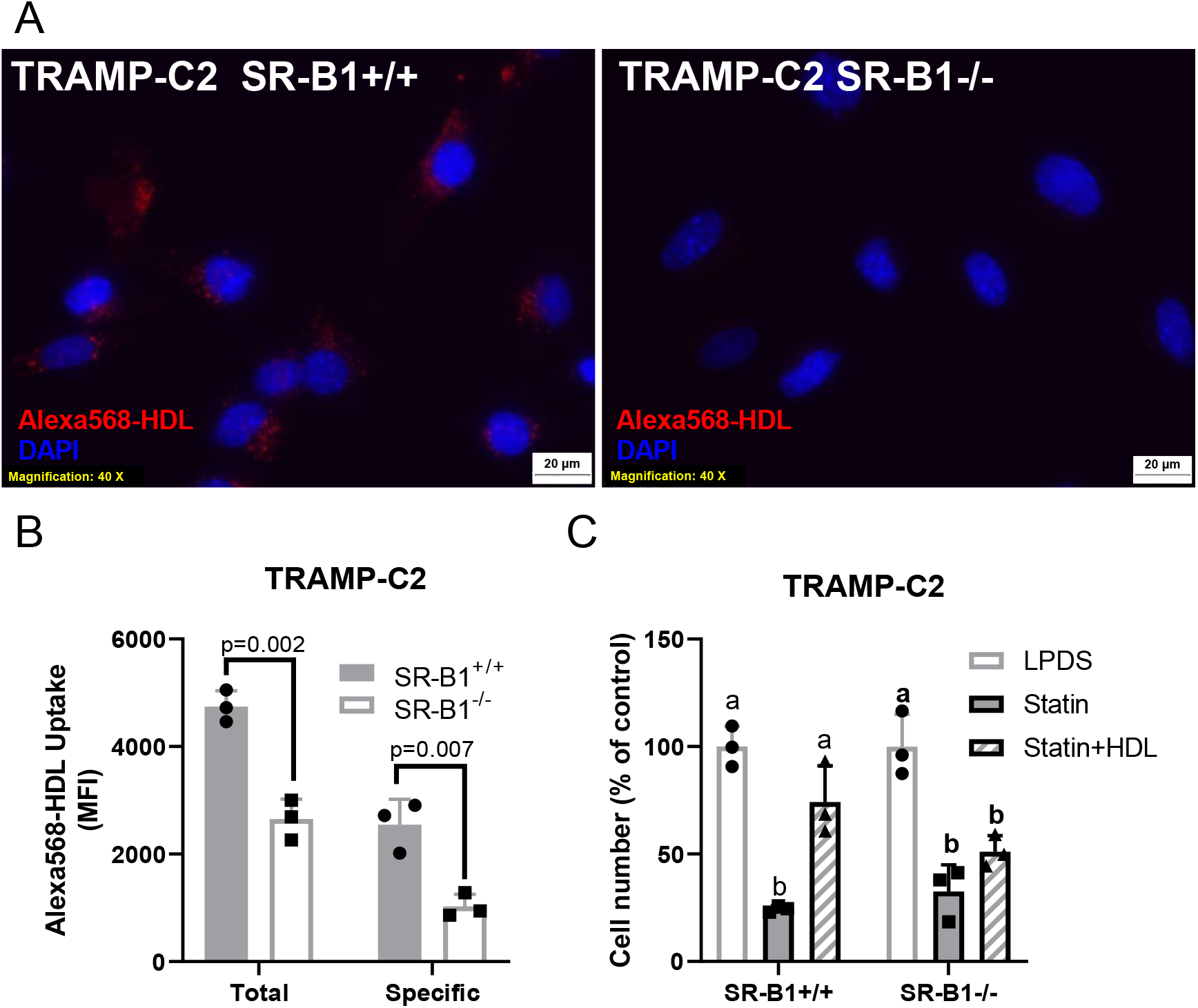
HDL uptake and impact on statin treated cells. TRAMP-C2 SR-B1^+/+^ or SR-B1^−/−^ cells were incubated with 20 μg/ml Alexa568-HDL for 90 minutes, then counter stained with DAPI. A) Epifluorescent microscopy. B) Total and specific uptake analysis by flow cytometry with TRAMP-C2 SR-B1^+/+^ and SR-B1^−/−^ cells treated with 20 μg/ml Alexa568-HDL and ± 2mg/ml unlabeled HDL for 90 minutes in serum free media (n=3, mean ± SD; t-test p-values displayed). C) Cell accumulation of TRAMP-C2 SR-B1^+/+^ and SR-B1^−/−^ in LPDS treated with ± 1 μM lovastatin and 100 μg/ml HDL for 4 days normalized to untreated cells (n=3, mean ± SD; different letters represent p<0.05 by ANOVA with Tukey posttest within each cell type).

### HDL and SR-B1 effects on prostate cancer progression in vivo

We hypothesized that elevated HDL levels may promote tumor progression in an SR-B1-dependent manner. To test our hypothesis, we utilized C57BL/6J WT and apoA1-KO mice as high and low HDL models. WT mice had fasting total and HDL cholesterol levels of 97±15 mg/dL and 63±17 mg/dL, respectively. apoA1-KO mice had significantly lower levels of total and HDL-cholesterol (32±8.mg/dL and 18±9 mg/dL, respectively) vs. WT hosts (p<0.0001 for both, Fig. S1A in the *Online Supporting Information*). WT mice weighed more than apoA1-KO mice (p<0.0001, Fig. S1B); however, there were no differences in testes weights (Fig. S1C). We measured plasma testosterone, and the data was not normally distributed with several outliers. Nonparametric T-tests found no effect on testosterone levels between WT and apA1-KO mice (Fig. S1D); however, removal of the two outliers in each group, resulted in normally distributed data showing 42% reduced testosterone levels in apoA1-KO mice (p=0.009, Fig. S1E). Dihydrotestosterone levels were undetectable in most mice (not shown).

To test our central hypothesis of the effects of host HDL and tumoral SR-B1 status on tumor progression, we performed a four-arm study using 2×10^6^ syngeneic TRAMP-C2 SR-B1^+/+^ or SR-B1^−/−^ cells that were subcutaneously injected into WT or apoA1-KO mice on the C57BL/6J background. Over an 8-week time course SR-B1^+/+^ and SR-B1^−/−^ cells formed solid tumors in WT and apoA1-KO mice. Histology of H&E stained tumors from all study arms showed unorganized sheets of cells with irregular shaped nuclei (Fig. S2). Tumors from all group were characterized as aggressive by a clinical pathologist. Tumor volume (p<0.0001, Fig. 6A) and survival (p=0.0016, Fig. 6B) were significantly different in the treatment arms.

**Figure 6.**
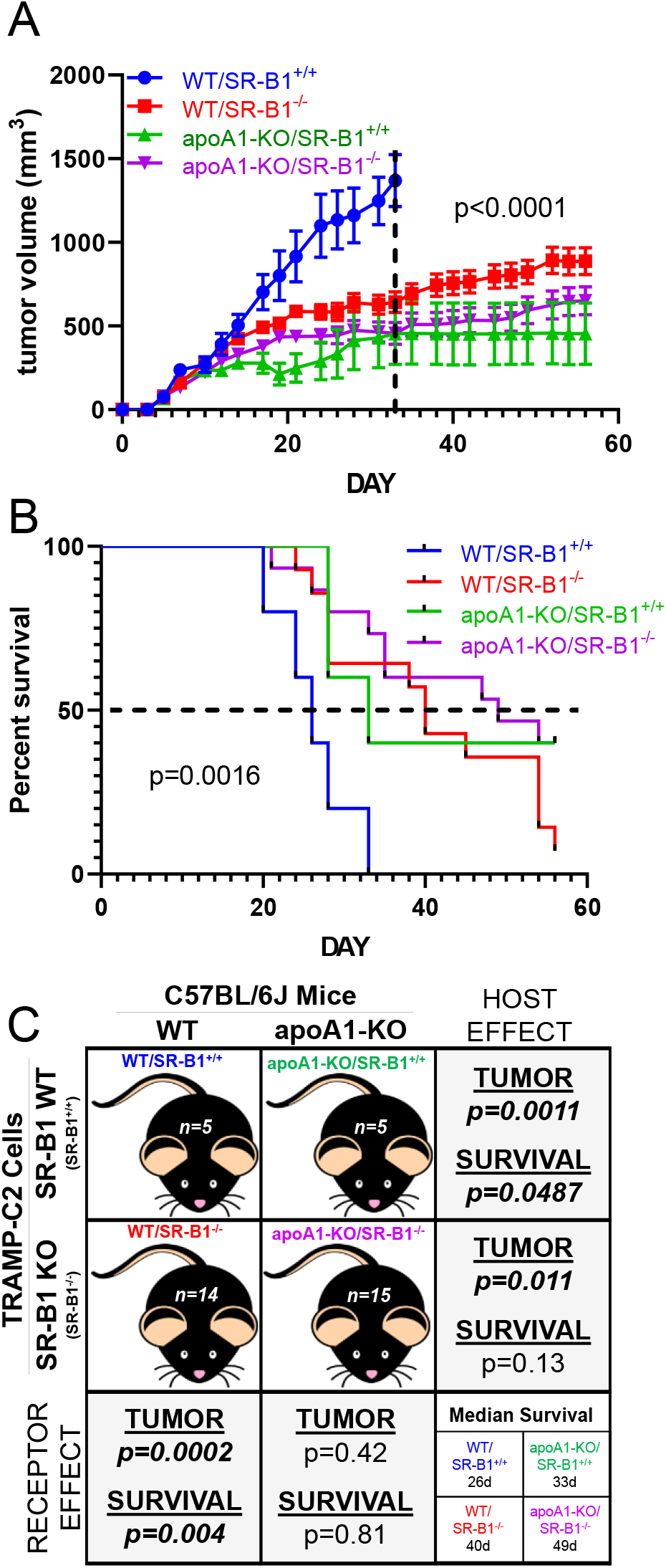
Tumor progression study *in vivo*. A) Tumor volumes of mice injected with 2×10^6^ cells in left and right flanks, then followed for 8 weeks. Tumor volume is expressed as mean ± SEM; WT mice SR-B1^+/+^ cells (WT/SR-B1^+/+^, blue) n=5; WT mice SR-B1^−/−^ cells (WT/SR-B1^−/−^, red) n=14; apoA1-KO mice SR-B1^+/+^ cells (apoA1-KO/SR-B1^+/+^, green) n=5; apoA1-KO mice SR-B1^−/−^ cells (apoA1-KO/SR-B1^−/−^, purple) n=15. Two-way ANOVA up to day 33 when all groups survived demonstrated group effects, time effects, and interaction effects all with p<0.0001. B) Kaplan Meier survival plot (p=0.0016 by Mantel Cox Log Rank Sum test for all four groups). C) Summary of study design and pairwise statistical analysis of host mouse genotype and injected cell genotype effects tumor progression and survival. Tumor volume analyses involving WT/SR-B1^+/+^ group, only used data up to day 33. Pairwise two-way ANOVA group effect on tumor volumes p-values are displayed. Pairwise survival Mantel Cox Log Rank Sum group effect p-values are displayed.

First examining the host mouse effects (shown in the rows in Fig. 6C) using SR-B1^+/+^ cells, WT mice injected with SR-B1^+/+^ cells (WT/SR-B1^+/+^) had significantly larger tumors vs. apoA1-KO mice injected with SR-B1^+/+^ cells (apoA1-KO/SR-B1^+/+^) (p<0.0001, Fig 6A, C). Additionally, the WT/SR-B1^+/+^ group all reached their human endpoint by day 33 with median survival of 26 days, and their log-rank survival was significantly shorter vs. the apoA1-KO/SR-B1^+/+^ group (median survival of 33 days, p=0.049, Fig 6B,C). Examining the host effects using SR-B1^−/−^ cells, WT mice injected with KO cells (WT/SR-B1^−/−^) had significantly larger tumors vs. apoA1-KO mice injected with SR-B1^−/−^ cells (apoA1-KO/SR-B1^−/−^) (p<0.0001, Fig 6A, C). The WT/ SR-B1^−/−^ group (median survival of 40 days) trended towards shorter log-rank survival vs. the apoA1-KO/SR-B1^−/−^ group (median survival of 49 days, p=0.13, Fig 6B, C). Thus the host effect was clear, mice with higher HDL levels promoted more rapid tumor progression, with shorter survival.

Next examining the SR-B1 receptor effects in WT hosts (shown in the columns in Fig. 6C), the WT/SR-B1^+/+^ group had significantly larger tumors vs. WT mice injected with SR-B1^−/−^ cells (WT/SR-B1^−/−^) (p<0.0001, Fig 6A, C). Log-rank survival for the WT/SR-B1^+/+^ group was significantly shorter vs. the WT/SR-B1^−/−^ group (p=0.004, Fig 6B, C). Examining the SR-B1 receptor effects in apoA1-KO mice, apoA1-KO mice injected with SR-B1^+/+^ cells (apoA1-KO/SR-B1^+/+^) had similar rates of tumor progression and survival compared to the apoA1-KO mice injected with SR-B1^−/−^ cells (apoA1-KO/SR-B1^−/−^) (NS, Fig 6A, B, C). Thus the SR-B1 receptor effect was only evident in WT mice, where receptor expression promoted more rapid tumor progression and shorter survival.

## Discussion

The effects of HDL are mixed on the prevalence of many cancers including, prostate, breast, endometrial, gynecologic, colorectal, biliary tract, lung, and hematological cancers (reviewed in (26)). Prostate cancer is a disease typically driven by androgens in which surgical and chemical castration have proven to slow progression of the disease (27,28). However, prostate cancer often becomes resistant to such treatments (29). The effects of HDL on prostate cancer development and progression are controversial, with both positive and inverse associations found in various studies (2–5). A meta-analysis of 14 large prospective studies found no significant effect of HDL-cholesterol on the risk of prostate cancer (30). Additionally, a Mendelian randomization study showed that the 35 SNPs associated with HDL-cholesterol did not generate a genetic risk score for prostate cancer (31). Thus, HDL levels may not affect the development of prostate cancer, but may still influence its progression.

HDL-cholesterol is sexually dimorphic, being ~10 mg/dl higher in adult women vs. men (32). However, before puberty, boys and girls have similar (high) HDL, which drop as boys go through puberty and stay unchanged as girls go through puberty (18,19). Androgens lower HDL in humans, and this is thought to be mediated by androgen induction of hepatic SR-B1(21). Thus, the effect of androgens on lowering HDL-cholesterol may partially obscure a potential positive association between HDL and prostate cancer, as androgens drive the early stages of prostate cancer.

HDL is synthesized from lipid-poor apoA1 by ABCA1; and, HDL cellular uptake is mediated by SR-B1 (8,9). Lee et al. found that ABCA1 expression was lower and ABCA1 promoter had higher DNA methylation in more vs. less advanced prostate cancer, and that ABCA1 gene expression was epigenetically silenced by DNA methylation in the LNCaP cell line (24). Schörghofer et al. demonstrated that SR-B1 mRNA expression was higher in high grade vs. low grade prostate cancer biopsies, and also higher in metastatic vs. primary prostate cancer (22). In contrast, this study reported no difference in LDLR expression in these tissues (22). Gordon et al. also reported increased SR-B1 mRNA expression, but lower LDLR mRNA expression, in prostate cancer vs. matched normal prostatic tissue (23). Our TCGA data analysis showed that SR-B1 mRNA is upregulated and that ABCA1 mRNA is decreased in prostate cancer vs. paired adjacent normal tissue, in agreement with the above studies (22,23).

We found that HDL treatment of cells cultured in LPDS led to an increase in cell number and proliferation in both a human and a mouse prostate cancer cell line. A similar growth promoting effect of HDL was observed by Sekine et al., using prostate cancer cell lines cultured in 1% FBS This HDL effect was associated with increased phospho-ERK1 and phospho-AKT after 30 minutes of HDL incubation (33). In both TRAMP-C2 and DU145 cells, we found that SR-B1 knockout abolished the growth promoting effects of HDL. However, Sekine et al. showed that siRNA-mediated knockdown of SR-B1 in PC3 cells did not abolish the growth promoting effects of HDL, which they attributed to ABCA1 expression (33). Gordon et al. reported that the growth of C4-2 human prostate cancer cell line in 10% FBS is inhibited by SR-B1 siRNA or by the anti-SR-B1 drug BLT-1, agreeing with our finding that SR-B1 is growth promoting in the presence of HDL.

Sekine et al. demonstrated that HDL promotes cholesterol uptake by PC3 prostate cancer cells, yet 100 μg/ml HDL in 1% FBS for 24h it did not promote an increase in total cholesterol levels in PC3, DU145, or LNCaP cell lines (33). However, we demonstrated that 200 μg/ml HDL in LPDS for 48h increased total cholesterol in DU145 and TRAMP-C2 cells (Fig. 2A, B). Furthermore, we showed that Alexa568-HDL uptake by TRAMP-C2 cells was in part mediated by SR-B1 (Fig. 5A, B), a finding that was demonstrated by Gordon et al., in C4-2 prostate cancer cells in which DiI-HDL uptake was partially reduced upon SR-B1 knock down or chemical inhibition by BLT-1 (23). We found that inhibiting *de novo* cholesterol synthesis in TRAMP-C2 cells by lovastatin led to reduced cell accumulation, which was rescued by HDL treatment in an SR-B1 dependent fashion, suggesting that the cholesterol content of HDL may play a role in the recovery of cell accumulation (Fig. 5C). Additionally, various statins were found to promote cell cycle arrest of PC3 cells, also indicating the need for *de novo* cholesterol biosynthesis to drive prostate cancer cell proliferation (34). Four meta-analyses of human observational studies on statin use and prostate cancer incidence, progression, and mortality have been published in 2016 or later, with three finding beneficial effects of statins (35–38). For example, prostate cancer specific mortality was significantly reduced in statin users pre- and post-diagnosis with prostate cancer (HR =0.53 and 0.64, respectively) (35). Thus, cholesterol metabolism may play an important role in prostate cancer therapeutics.

We have previously shown that phosphatidylinositol 4,5-bisphosphate (PIP2) can be trafficked in and out of cells via SR-B1 and ABCA1 and that the majority of circulating PIP2 is carried on HDL(39). PIP2 is a phospholipid primarily sequestered on the inner leaflet of the plasma membrane, where it can be converted to PIP3 with subsequent recruitment and activation of AKT (Reviewed in (40)). Thus, HDL-PIP2 uptake via SR-B1 into prostate cancer cells may also promote cell proliferation. Semenas et al. showed that inhibition of PI-4-phosphate 5 kinase 1 alpha (PIP5K1α), which produces PIP2, reduced prostate cancer tumor growth, androgen receptor (AR) expression, and prostate specific antigen (PSA) levels (41).

A large meta-analysis demonstrated that increased HDL-cholesterol was associated with lower incidence of cancer; although this study did not examine different types of cancer (42). This finding was corroborated in C57BL/6J mice using syngeneic B16F10 melanoma and Lewis Lung carcinoma cells injected into (going from low to high HDL-cholesterol levels) apoA1-KO, WT, and human apoA1 transgenic mice; and, for both of these cancer types higher HDL leads to smaller tumors (43). The protective effect of HDL was associated with increased tumor associated macrophages, cytotoxic CD8 T-cells, and decreased recruitment of myeloid derived suppressor cells (43,44). In addition, B16F10 melanoma tumor progression was slower in *Scarb1* KO mice, another model of high plasma HDL (44). In C57BL/6J WT and apoA1-KO mice, we found an opposite effect of host HDL on syngeneic TRAMP-C2 prostate cancer cells, where higher HDL led to larger tumors, and decreased survival.

Why do the TRAMP-C2 cells respond differently than the B16F10 and Lewis Lung cells? Perhaps it is due to the relative expression levels of SR-B1 and the role of HDL in providing lipids required for cell cycling. Llaverias et al., reported that feeding a western-type vs. chow diet to C57BL/6J TRAMP transgenic mice, which are prone to spontaneously develop prostate tumors, results in increased total and HDL-cholesterol, and a higher prostatic tumor incidence with larger tumors (45). The tumors from western type vs. chow diet-fed fed TRAMP mice also have higher levels of SR-B1 expression (45). The SNP rs4765623, intronic in the *SCARB1* gene, is associated with clear cell renal cell carcinoma (ccRCC), a cancer characterized by excessive lipid loading (46). This same SNP is associated with *SCARB1* expression levels in human left ventricle, with the T allele associated with increased expression (47). rs4765623 is in linkage disequilibrium with rs12582221(d’=1, r2=0.39) (48), and this SNP is highly associated with SR-B1 expression in testes (47). SR-B1 expression is increased in ccRCC tissue vs normal kidney tissue (49,50), similar to what we and others observed in prostate cancer vs. normal prostate. Also in alignment with our SR-B1 KO findings in prostate cancer cells, Velagapudi et al., showed that antibodies against SR-B1 reduce cellular uptake of ^125^I-HDL into, and HDL-induced proliferation of, a ccRCC cells line (51). Thus, SR-B1 and HDL may drive proliferation of other lipid accumulating cancers. Whether HDL promotes or inhibits tumor progression may depend on tumor lipid delivery vs. host immune effects.

Our analysis and two other studies found higher SR-B1 mRNA levels in prostate cancer vs. normal prostate tissue, in high grade vs. low grade prostate cancer, and in metastatic vs. primary prostate cancer (22,23). Some prostate cancer cells may be able take up HDL-cholesterol via SR-B1 to synthesize *de novo* androgens (23,52). In general, SR-B1 expression is highest in the major HDL metabolizing organ, the liver, and in steroidogenic tissues. In humans SR-B1 expression is highest in adrenals, liver, and ovary (47). In 9 tested tissues of C57BL/6J mice, the liver and testes had the highest expression of SR-B1 (53). The role of HDL uptake to support CE storage for steroidogenic tissue was demonstrated in apoA1-KO mice, where adrenal and ovarian CE stores, as wells as stress-induced plasma corticosteroid levels, are significantly reduced (54). Although total testes CE storage was not different between WT and apoA1-KO mice, histological analysis showed markedly reduced neutral lipid loading in the leydig cells, the site of *de novo* androgen biosynthesis (54). We measured plasma testosterone levels in 2 cages each of WT and apoA1-KO mice, and in each cage, there was one high outlier, which may be due to the well-known effect of higher testosterone levels in the socially dominant male of group-housed mice (55,56). Excluding these outliers, we found 42% higher plasma testosterone in the WT vs. apoA1-KO mice, which is a confounder for the pro-tumor effect of HDL in our study.

We found that SR-B1^−/−^ vs. SR-B1^+/+^ TRAMP-C2 cells injected into WT hosts with high HDL-cholesterol, led to reduced tumor progression and increased survival; thus, SR-B1 promotes prostate cancer tumor growth *in vivo*, similar to SR-B1’s growth-promoting effects we observed in cell culture. This is similar to the finding of Gordon et al., who showed that treatment of mice with the SR-B1 inhibitor BLT-1 led to reduced human PC3 cell xenograft progression (23). These findings align well with our TGCA analysis and prior analyses showing that SR-B1 expression is higher in prostate cancer in more advanced prostate cancer vs. controls (22,23). Of interest, we also noted increased tumor progression of SR-B1^−/−^ TRAMP-C2 cells in the WT vs. apoA1-KO hosts, which we attribute to the nonspecific uptake of HDL into the SR-B1^−/−^ cells *in vivo*, which we observed in these cells in tissue culture (Fig. 5B). It appears that the growth promoting effects of SR-B1 in TRAMP-C2 cells is entirely mediated by its ligand HDL, as SR-B1 status (KO vs. WT) has no effect on tumor progression in apoA1-KO mice.

In conclusion, our findings are in alignment in with Llaverias et al., where a high fat diet increased autochthonous tumor prevalence and burden in the TRAMP transgenic mouse model, associated with increased HDL levels (45). Additionally, our findings corroborated the *in vivo* findings of Gordon et al., who demonstrated that treatment with an SR-B1 inhibitor decreased prostate cancer progression (23). Our study was more specific and could differentiate cancer cell vs. host effects due to genetic ablation of SR-B1 in the cancer cells, whereas Gordon et al. used a chemical inhibitor of SR-B1, having specific and non-specific effects on the host as well as the xenograft cancer cells. Our study employed immunocompetent mice, which may be a better model to study HDL effects on prostate cancer since HDL has been shown to modulate tumor infiltrating leukocytes (43).

## Experimental procedures

The human prostate cancer cell line DU145 and mouse prostate cancer cell line TRAMP-C2 were purchased from American Type Culture Collection (Manassas, VA). DU145 cell line was authenticated by Genetica cell line testing. DU145 was cultured in RPMI1640 (supplemented with 10 % FBS (Sigma) and TRAMP-C2 was cultured in DMEM supplemented with 10% FBS (Sigma) and 10 nM dihydrotestosterone (Sigma) at 37 °C in 5 % CO_2_. Both cell lines grown treated with low-does mycoplasma removal agent (MP Biomedicals) as a preventative measure. Mycoplasma testing was performed regularly to confirm absence of mycoplasma contamination on cells with SR-B1 rabbit polyclonal antibody (NB400-104) was purchased from Novus Biologicals. GAPDH rabbit polyclonal antibody (sc-25778) was purchased from Santa Cruz. Lipoprotein deficient serum (LPDS) was prepared as previously described (57). In brief, FBS was adjusted to 1.21 g/ml, then centrifuged to separate lipoprotein from sera. LPDS was collected and dialyzed against PBS (pH 7.4), and then sterilized using a 0.22 μm filter. LPDS was adjusted to 30 mg/ml protein using PBS.

For Lipoportein isolation and labeling expired de-identified human plasma was collected from normal healthy volunteers was acquired from the Cleveland Clinic blood bank under an exempt Institutional Review Board protocol. LDL (1.019–1.063 g/mL) and HDL (1.063–1.21 g/mL) were isolated by potassium bromide density gradient centrifugation as described (57). Lipoproteins were dialyzed against PBS (pH 7.4), sterilized using a 0.22 μm filter, and then protein concentration was determined by the Alkaline-Lowry method (58). HDL was labeled with Alexa568 by incubating 4.5 mg Human HDL in 90 μl 1 M sodium bicarbonate with Alexa Fluor 568 succinimidyl ester (A20003, Thermo Fisher) for 1 hr at room temperature (8:1 dye:estimated apoA1 mole ratio). The reaction was stopped by incubating the conjugate with 0.1 ml of 1.5 M hydroxylamine (pH 8.5) for 1 hr at room temperature. The conjugate was purified by extensive dialysis with PBS. For murine lipoprotein isolation and quantification, blood from the retro-orbital venous sinus was collected and centrifuged at 12,000xg for 30 minutes.

For cell accumulation, proliferation and cell cycle analysis, cells were seeded into a 24-well at densities of 10,000 and 20,000 cells/well for TRAMP-C2 and DU145 cells, respectively in 10% FBS containing media overnight, then incubated with serum free media for 30 minutes. Cells were then incubated in 10% LPDS with or without 300 μg/ml HDL in the absence or presence of 1 μM Lovastatin for 4 days. Thereafter, cells were lifted by trypsin and counted using the automated cell counter (Z series) by Beckman Coulter. For cell proliferation analysis, cells were labeled with 2 × 10^−6^ M PKH26 (Sigma) for 5 minutes, then plated in 10% FBS media overnight and subsequently treated with 300 μg/ml HDL in 10% LPDS media for 5 days and quantified by flow cytometry. For cell cycle analysis, cells were treated with or without 300 μg/ml HDL in 2% LPDS media for 1 day, ethanol fixed, stained with 50 μg/ml propidium iodide, and then subjected to flow cytometry.

In order to generate SR-B1 KO cell lines, TRAMP-C2 cell were transfected with the Cas9 expression plasmid, pSpCas9(BB)-2A-Puro (Addgene #48139) using Lipofectamine LTX & Plus Reagent (Invitrogen) according to manufacturer instructions, then treated with 5 μg/ml puromycin for 2 days to clonally select for Cas9 stably transfected TRAMP-C2 cells. Nucleofection (Amaxa), according to manufacturer, was used to transfect 1×10^6^ Cas9 stable TRAMP-C2 cells with 0.6 nM mouse SR-B1 sgRNA targeting exon 4. Human DU145 cells (1×10^6^) were co-transfected with 0.6 nM human SR-B1 sgRNA targeting exon 4 complexed with 0.07 nM Cas9 protein (Synthego) by nucleofection. Both sgRNAs were designed using CRISPOR online software (59) and synthesized by Synthego. sgRNAs sequences: mouse sgScarb1 reverse strand 5’-gguccacgcucccggacuac -3’; human sgSCARB1 reverse strand 5’- caugaaggcacguucgccga-3’. Transfected cells were plated in 96-well dishes to approximately 1 cell/well then clonally expanded and screened via PCR-Sanger sequencing to detect targeted sequence disruptions, and then by Western blot for SR-B1. PCR screening primers: human fwd 5’-ccagtgggttctgagtttccca-3’, rev 5’-gatccccagccagctacaaagc-3’; mouse fwd 5’-ggttccatttaggcctcaggt-3’, rev 5’-ctctctgaagggacagaagacac-3’.

For HDL uptake assays, approximately 100,000 cells/well were seeded onto a 24-well plate and incubated overnight in 10% FBS containing media. Cells were then incubated with serum free media for 30 minutes and then treated with 20 μg/ml Alexa568-HDL, as described above, for 90 minutes. Cells were fixed, then counterstained with DAPI for microscopy, or stained first, then formalin fixed for flow cytometry.

To evaluate cholesterol mass, cells were plated on a 6-well plate and incubated overnight in the medium, 10% FBS containing media. Cells were incubated in serum free media for 30 minutes and treated with or 200 μg/ml HDL for 2 days. Cholesterol concentrations were measured through by an enzymatic fluorescent assay and normalized to protein as described by (60).

For protein expression via Western blot, cell lysates were prepared in RIPA buffer (Pierce), containing 1 mM phenylmethylsulfonyl fluoride and 10% protease inhibitor cocktail (Sigma). Equal amounts of proteins were electrophoresed on 4–20% SDS–PAGE and transferred to polyvinylidene difluoride membranes. Each membrane was incubated with a 1:2000 dilution of primary antibodies described above. Blots were developed with a 1:5000 dilution of the HRP-conjugated secondary antibody (BioRad). Proteins were visualized, using Amersham Hyperfilm ECL (GE Healthcare).

Our in vivo studies used 20–22-Week-old, age-matched, apoA1-KO on and WT C57BL/6J male mice were subcutaneously injected in both flanks with 2×10^6^ SR-B1 KO or WT TRAMP-C2 (syngeneic) prostate cancer tumor cells per site. Tumor progression and body weights were assessed three times per week for eight weeks or until reaching an experimental endpoint of tumor reaching 15 mm in diameter, tumor ulcerations, impaired mobility, or 20% loss in body weight. Tumor volume, based on caliper measurements, was calculated according to the ellipsoid volume formula, tumor volume = (the shortest diameter)^2^ × the largest diameter × 0.525. After a mouse reached its endpoint, its final tumor volumes were included in the analysis up to the day that all mice from that group reached its endpoint, after which data were no longer plotted. All experiments and procedures were approved by Institutional Animal Care and Use Committee (IACUC) of the Cleveland Clinic, Cleveland, OH, USA (Protocol No. 2016-1722).

Bioinformatic information was acquired and viewed by The UCSC Xena browser (61)was used to visualize and extract expression data for *SCARB1*, *LDLR*, and *ABCA1* in paired normal adjacent and prostate cancer tissue from the TCGA PRAD dataset. The Genotype-Tissue Expression (GTEx) Project (47) was supported by the Common Fund of the Office of the Director of the National Institutes of Health, and by NCI, NHGRI, NHLBI, NIDA, NIMH, and NINDS. The data used for the analyses described in this manuscript were obtained from the following independent searches [SCARB1], [rs4765623], [rs12582221], on the 0424.v8.p2. LDlink (48) was used to evaluate the linkage disequilibrium of *SCARB1* SNPs [rs12582221] and [rs4765623] in the CEU population of the 1000 Genomes Project, with reference genome GRCh37/gh19 on 01/08/2020; LDlink phase 3, version 5.

Statistical analysis of TGCA PRAD gene expression data from matched pairs of prostate cancer and normal adjacent tissue was analyzed by the Wilcoxon paired non-parametric signed rank test, and % changes were calculated from the antilog 2 of the median values. All *in vitro* data are expressed as the mean ± SD of at least triplicates. Differences between the values were evaluated by either the Student T-test, with Bonferroni correction for multiple tests when appropriate, or one-way analysis of variance (one-way ANOVA) with Tukey’s or Dunnett’s post-hoc analysis, with p <0.05 considered statistically significant. ANOVA annotation uses lettering (a, b, c, etc.), where groups not sharing the same letter indicates p<0.05. All *in vivo* tumor volume data is expressed as mean ± SEM Differences between the tumor volumes during the time course were evaluated two-way analysis of variance (two-way ANOVA). Survival curve data was evaluated by Mantel-Cox Log Rank analysis. Statistics were performed using GraphPad Prism V.8.1.1 software.

## Acknowledgements

We would like to thank the following cores at the Cleveland Clinic Lerner Research Institute: The Biological Research Unit (BRU) for assistance with mouse handling, the Flow Cytometry Core for advice and help with flow cytometry instrumentation, and the Histology Core for tissue staining. We would also like to thank Dr. Erinn Downs Kelly for comments on tumor pathology. Lastly, we would like to acknowledge the funding received from the Case Comprehensive Cancer Center pilot grant program funded by NCI grant P30CA043703.

## FOOTNOTES

Funding for this research and training was suppored by: The National Institutes of Health grants R01HL128628 (J.D.S), R01DK120679 (J.M.B), P50AA024333 (J.M.B), P01HL147823 (J.M.B), R01CA172382 (N.S), R01CA190289 (N.S), R01CA236780 (N.S.), F31HL134231 (C.A.T), an American Heart Association grant SDG25710128 (K.G.), the Case Comprehensive Cancer Center pilot grant RES511106 (J.D.S), and the VeloSano Foundation (J.D.S). Disclaimer: The content is solely the responsibility of the authors and does not necessarily represent the official views of the National Institutes of Health.

## The abbreviations used are

apoA1: apolipoprotein-AI
BLT-1: Block Lipid Transport-1
ccRCC: clear cell renal cell carcinoma
CE: Cholesterol ester
HDL: High-density Lipoprotein
KO: knockout
LDL: Low-density Lipoprotein
LDLR: Low-density Lipoprotein Receptor
LPDS: Lipoprotein Deficient Serum
MFI: Median Fluorescence Intensity
PIP2: phosphatidylinositol 4,5-bisphosphate
PIP5K1α: Phosphatidylinositol-4-Phosphate 5-Kinase Type 1 Alpha
PRAD: Prostate Adenocarcinoma
PTEN: Phosphatase and tensin homolog
SR-B1: Scavenger Receptor Class B, Type 1
sgRNA: single guide RNA
TCGA: The Cancer Genome Atlas
TRAMP: Transgenic Adenocarcinoma Mouse Prostate

